# Glycerol alone effects 1,3-propanediol production via the aerobic propanediol utilization pathway in *Salmonella enterica*

**DOI:** 10.64898/2026.03.20.713204

**Authors:** Madeline R. Joseph, Brett J. Palmero, Nolan W. Kennedy, Danielle Tullman-Ercek

## Abstract

Crude glycerol is an underutilized waste stream. Viable routes for converting it to 1,3-propanediol (1,3-PDO) can conserve important resources and add value to its supply chain. Biological methods are appealing because they can circumvent expensive preprocessing steps while operating under mild conditions. Here, we show that the propanediol utilization pathway of *Salmonella enterica* serovar Typhimurium LT2 can be used to convert glycerol, including unprocessed crude glycerol, into 1,3-PDO under aerobic conditions in minimal media. Additionally, we demonstrate that high concentrations of expensive cofactors are not necessary to achieve optimal production titers. This study lays the groundwork for continual iteration on this pathway for bioprocess development.

**Key points:**

- *S. enterica* can produce 1,3-propanediol from crude glycerol alone
- Glycerol-to-1,3-propanediol conversion is dependent on expression of the propanediol utilization (Pdu) pathway
- Sub-saturating concentrations of exogenous vitamin B_12_ can boost cell growth and 1,3-propanediol yield

## Introduction

Crude glycerol is an abundant and inexpensive byproduct of biodiesel manufacturing^1^. Although high-purity glycerol has several industrial applications, separating it from biodiesel side streams is economically and energetically inefficient^1,2^. The large supply and low price of crude glycerol, combined with the barriers to using it in traditional glycerol applications, make it a promising candidate as a substrate for high-volume chemical manufacturing^2^. Moreover, valorization of crude glycerol has the environmental benefits of avoiding landfilling, incineration, and ecological contamination, as well as displacing virgin feedstocks for bulk chemicals, particularly those derived from petrochemicals^1,3^.

One such platform chemical that could be manufactured from crude glycerol is 1,3-propanediol (1,3-PDO)^4,5^. This chemical has a large annual global demand (∼700 million USD) and a wide variety of application areas, particularly synthetic fiber production^3,6^. New products derived from1,3-PDO are in continuous development and may compete with traditionally petrochemical-derived products, such as polyurethane materials^6^. An efficient method for converting crude glycerol to 1,3-PDO would simultaneously add value to biodiesel side streams, conserve resources, and displace demand for fossil-based goods^7^.

Still, an optimal conversion strategy is not yet clear. Nonnegligible fractions of ash, salt, and fatty acids in most crude glycerol streams are known to deactivate catalysts, making chemical conversion challenging^1,8^. However, organisms that have evolved to transform contaminated, heterogenous carbon sources are less susceptible to these impurities. Employing biological chasses to facilitate the glycerol-to-1,3-PDO conversion thus presents an opportunity to overcome this roadblock and circumvent the need for intensive preprocessing^8^. There are several promising bacterial strategies but mostly fall short of industrial benchmarks^8,9^.

The 1,2-propanediol utilization (Pdu) pathway has emerged as a strong target for biological production of 1,3-PDO, but in limited organisms thus far^10–15^. Some species, notably *Lactobacillus reuteri* and *Klebsiella pneumoniae*, are known to use the Pdu pathway to convert glycerol to 1,3-PDO via an intermediate, 3-hydroxypropionaldehyde^11–14^. Genetic engineering of the *pdu* operon, which governs this pathway, has led to substantial increases in 1,3-PDO titers in *L. reuteri*, but the organism’s industrial potential is limited by its inability to grow on glycerol as a sole carbon source^11,13^. *K. pneumoniae* is an established 1,3-PDO production chassis, but the Pdu pathway plays a relatively minor role in 1,3-PDO production in this species^14^.

The Pdu pathway is more extensively studied in *Salmonella enterica*, in which 1,2-propanediol (1,2-PDO) is converted first to 1-propionaldehyde and then to either 1-propanol or propionate, which is used in central metabolism^16–22^. Despite greater characterization of the Pdu pathway in *S. enterica*, conversion of glycerol to 1,3-PDO through Pdu enzymes has not been investigated in this organism. Indeed, this reaction is barely remarked upon in literature at all, possibly because the Pdu pathway was not thought to be induced by glycerol robustly^17,19,23^.

In this work, we show for the first time that *S. enterica* can aerobically produce 1,3-PDO with glycerol as the sole carbon and energy source via the Pdu pathway. Moreover, we demonstrate that minimally preprocessed crude glycerol obtained directly from biodiesel manufacturing is a viable carbon and energy source for *S. enterica* to produce 1,3-PDO. Finally, we show that saturating doses of expensive cofactors are not necessary for optimal 1,3-PDO production *in vivo*. This work provides a preliminary basis for integrating the Pdu pathway into the development of economically viable, scalable systems for transforming crude glycerol to 1,3-PDO in aerobic bacterial chasses.

## Materials and Methods

### Strains

In all cases in this work, “wild-type” refers to *Salmonella enterica* serovar Typhimurium LT2. We produced the two genomically modified strains, Δ*pocR* and Δ*pduP*, from wild-type via the lambda red recombineering system, as described elsewhere^24,25^. All strains were grown from 45% glycerol stocks stored at -80 °C.

### Culture conditions

First, we streaked glycerol stocks onto agar plates supplemented with lysogeny broth Lennox (Dot Scientific DSL24066-500) and incubated them at 30 °C for 18-24 hours. We then isolated single colonies from these plates for overnight test tube cultures. These starter cultures grew in 5 mL lysogeny broth Miller (Fisher Scientific BP1426500) at 30 °C, 225 rpm, for 16 h.

For all experiments, we diluted the starter cultures 1:1000 in no-carbon essential (NCE) media (29 mM potassium phosphate monobasic, 34 mM potassium phosphate dibasic, 17 mM sodium ammonium hydrogen phosphate, 1 mM magnesium sulfate, and 50 µm ferric citrate) supplemented with cofactor adenosylcobalamin (AdoB_12_) and either pure or crude glycerol as the sole carbon source^24,26^. Unless otherwise noted, we supplemented all cultures with 55 mM glycerol and 150 nM AdoB_12_ (saturating concentration). These cultures grew at 37 °C, 225 rpm, over 24 h either in 500-mL glass Erlenmeyer flasks with a 100 mL working volume (Figs 1 and 2) or in 24-well polypropylene blocks (Corning P-DW-10ML-24-C) with a 5 mL working volume (Figs 3 and 2S).

**Figure 1.**
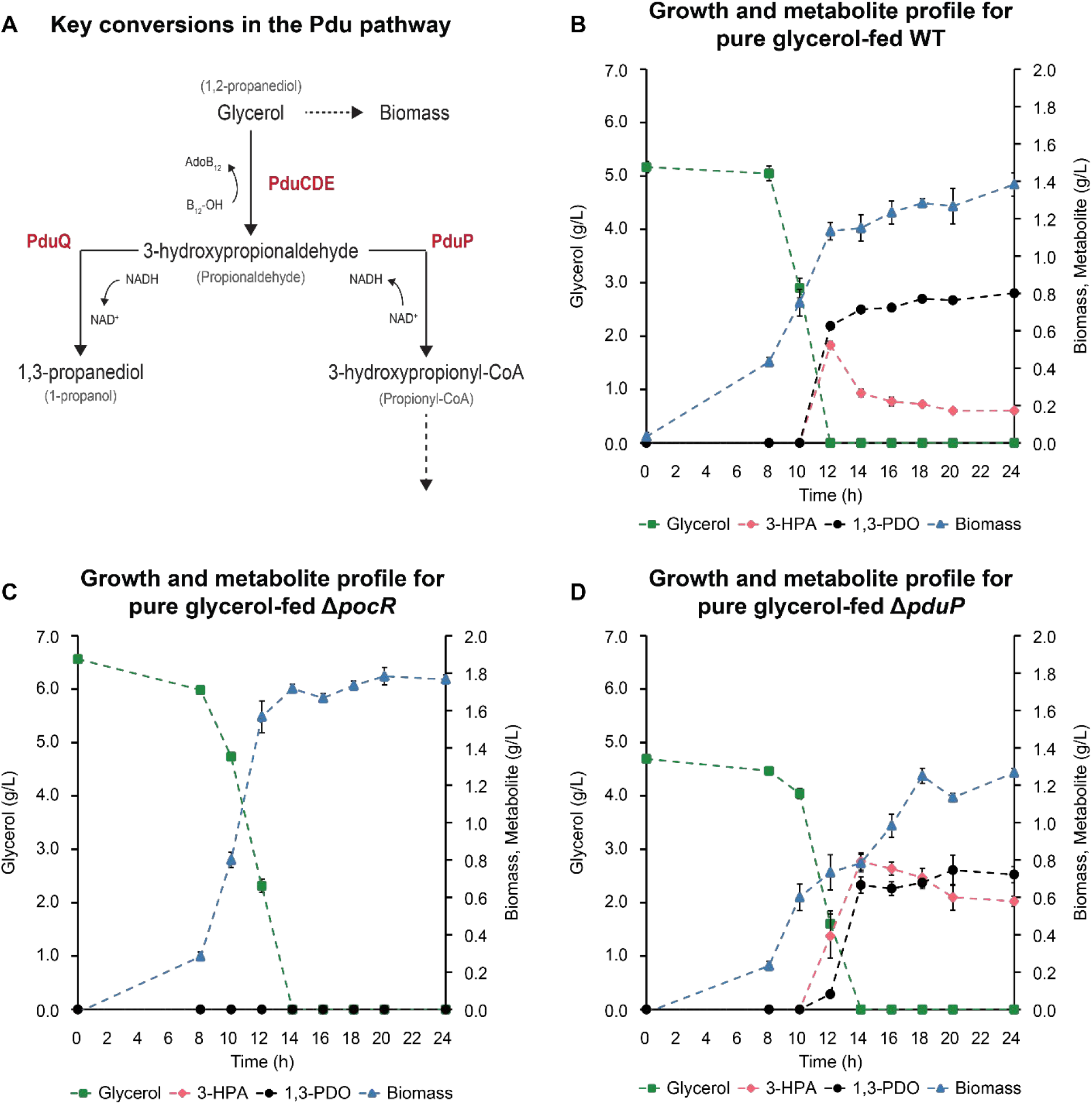
**A)** The 1,2-propanediol utilization (Pdu) pathway for glycerol-to-1,3-PDO conversion in *S. enterica*. Bold, red font indicates enzymes and small, grey parentheticals indicate metabolites in the 1,2-propanediol-induced system. Dashed lines represent conversions beyond the scope of this work. Progression of biomass accumulation, laboratory-grade glycerol consumption, and key metabolite production in **B)** WT *S. enterica*, **C)** a *pocR* knockout strain, and **D)** a *pduP*-deficient strain.

**Figure 2.**
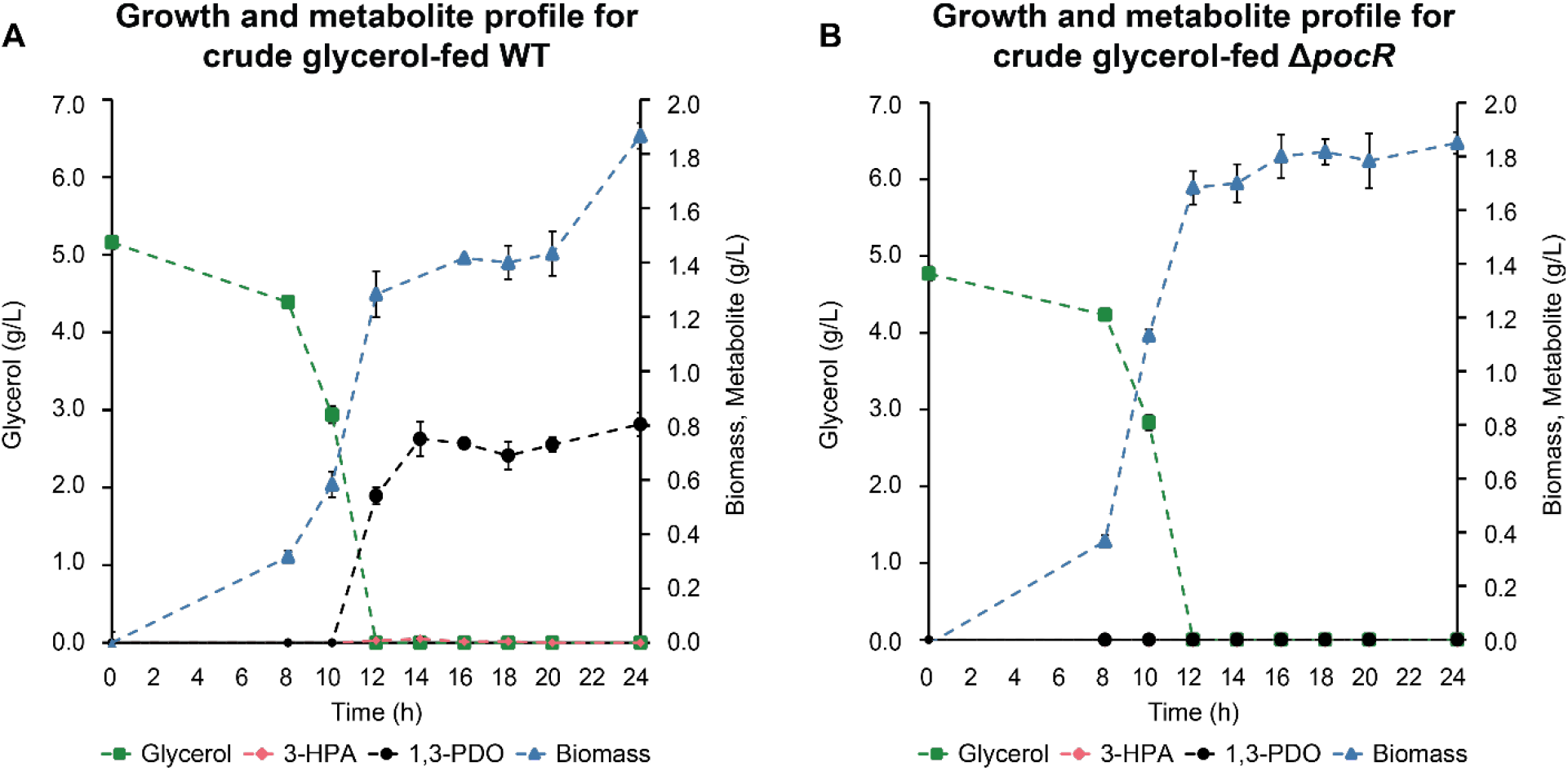
Progression of biomass accumulation, crude glycerol consumption, and key metabolite production in the **A)** wild-type and **B)** pocR-deficient strains of *S. enterica*.

**Figure 3.**
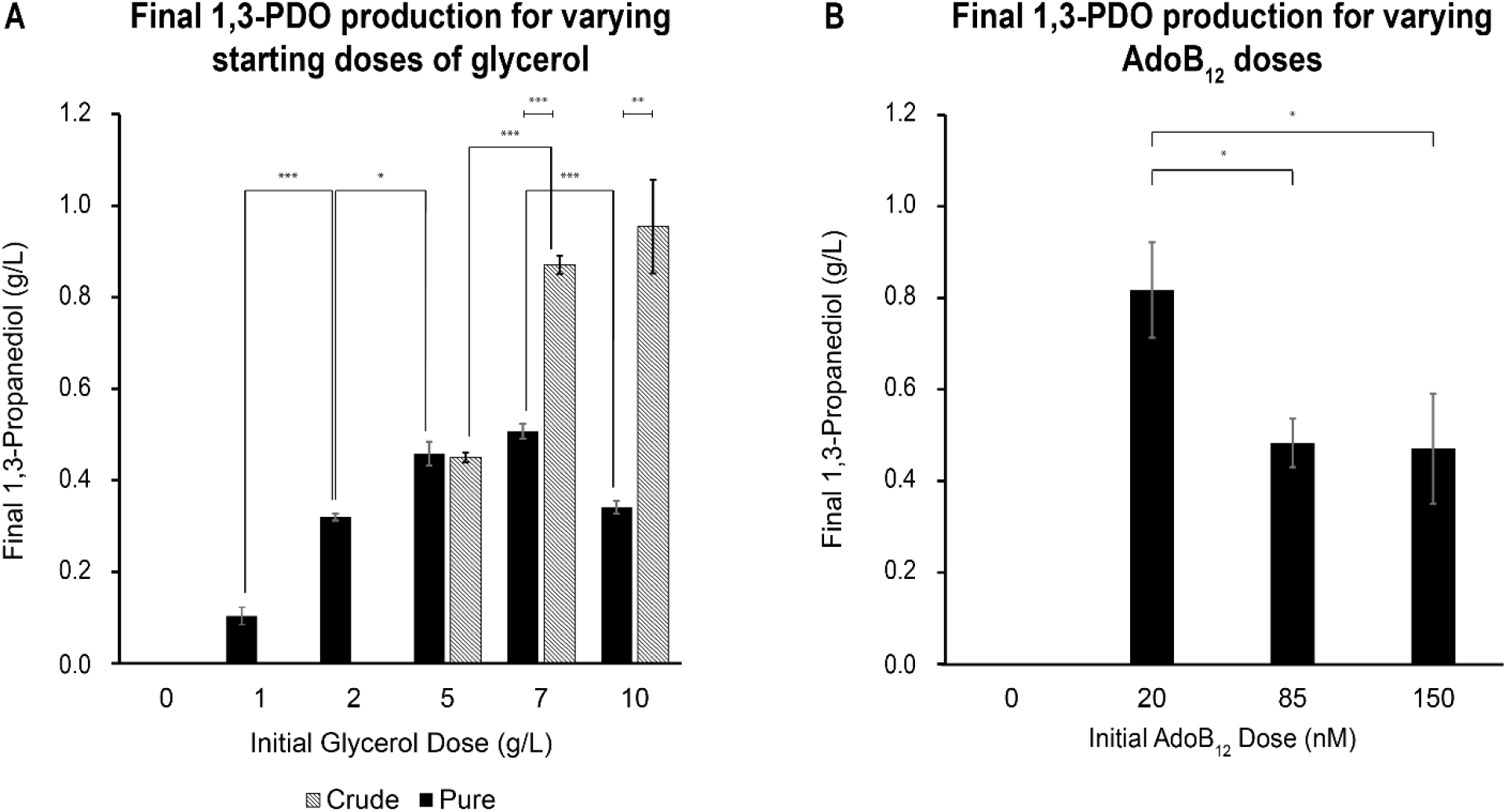
In wild-type *S. enterica*, 1,3-propanediol production at 24 h: **A)** for varying starting concentrations of laboratory-grade and crude glycerol. B) for varying doses (saturating, midpoint, limiting, and none, from left to right) of AdoB_12_. Asterisks denote p-value significance (*: *p* <0.05, **: *p* < 0.01, ***: *p* < 0.005).

We purchased >99% pure glycerol from Thermo Fisher Scientific (Fisher Scientific BP229-1). The Searle Biodiesel Lab at Loyola University Chicago, which manufactures biodiesel from various waste cooking oil streams, generously provided crude glycerol samples with methanol removal being the only preprocessing step. While the exact composition is unknown, we used high performance liquid chromatography (HPLC) to determine a glycerol fraction of approximately 50%.

### Biomass Measurements

All biomass growth is reported in g/L. When possible, optical density measurements at 600 nm (OD_600_) were preferred to increase throughput. We measured OD_600_ in a BioTek Synergy H1 microplate reader (Agilent 11-120-533). Samples were all processed in clear, flat-bottom 96-well plates (Greiner Bio-One 655101). If the culture’s OD_600_ appeared to exceed 1, a 10 µL sample was diluted in 90 µL uninoculated media before measuring OD_600_; otherwise, 100 µL of culture was measured directly. For each timepoint sampled, a new blank consisting of 100 µL uninoculated media was used. We determined conversion factors for OD_600_ to dry cell mass in the strains used in this work to report biomass in g/L (Fig S1)^27^.

Given the turbidity of crude glycerol, it was not possible in many cases to obtain biomass concentrations from OD_600_ measurements. Under these conditions, we measured dry cell mass directly^28^. For each timepoint, 2 mL of culture was transferred to a sterile 2-mL microcentrifuge tube and spun at 20,000 x *g* in a microcentrifuge (Eppendorf Centrifuge 5424) for 2 minutes. After discarding the supernatant, we washed and resuspended the cell pellets in NCE. The centrifugation and wash steps were repeated twice. We then allowed the cell pellets to dry in a 95 °C dry heat bath for 48-72 h (no mass fluctuation). After cooling, we weighed the cells in their respective premeasured tubes in an analytical balance (Ohaus EX124) to report dry cell mass in g/L.

### Metabolite Measurements

We used HPLC with protocols adapted from Palmero and collaborators (2025) to determine all metabolite concentrations^29,30^. Briefly, we sampled 100 µL of culture for each timepoint and centrifuged it for 5 minutes at 13,000 x *g* to pellet out cell debris. We then transferred the resulting supernatant to a filter tube (Corning 8169) and centrifuged again for 5 minutes at 13,000 x *g* to remove any residual impurities. Any samples not immediately analyzed were stored at -80 °C. Before running HPLC, we loaded 30 µL of thawed, well-mixed samples into 96-well plates (Bio-Rad HSP9601) with a 20 µL layer of mineral oil on top to prevent volatile compounds from evaporating.

Our Agilent 1260 Infinity II LC System injected 20 µL samples (Agilent G7111B and G7167A), which were separated by a Rezex™ ROA-Organic Acid H+ (8%) LC 670 Column (Phenomenex 00H-0138-K0) outfitted with a guard column (Phenomenex AJ0-8768). Column temperature was maintained at 35 °C (Agilent G7116A). We used 35% acetonitrile with 0.5 mM sulfuric acid as the mobile phase with a flowrate of 0.5 mL/minute over 30-minute intervals. A refractive index detector (RID) was used to monitor metabolites (Agilent G7162A). OpenLab ChemStation software performed peak integration on RID signals, which we converted to metabolite concentrations using commercially available standards of known concentration for glycerol (Fisher Scientific BP229-1), 3-hydroxypropionaldehyde (Ambeed Inc. A949844-1g), and 1,3-PDO (Sigma-Aldrich P50404-100G).

### Statistics

We performed all time course experiments in biological triplicate. In all figures, error bars represent one standard deviation in each direction based on the variation within these biological replicates. Significance was calculated via Welch’s Two-Sample t-test (Table S1).

## Results and Discussion

The primary native function of the Pdu pathway in *S. enterica* is to generate ATP and carbon compounds for central metabolism via 1,2-PDO metabolism^18–22^. The operon is activated by a positive transcriptional regulator (PocR) and encodes a metabolic cascade effected by several enzymes (PduCDELPQW)^17,31^. Though 1,2-PDO is the native inducer, glycerol can also induce the *pdu* operon in *S. enterica* under some conditions^17,19,23^. However, studies of glycerol induction are limited, and no resulting metabolite data have been documented in the literature for *S. enterica*. We present them here, showing that glycerol-to-1,3-PDO conversion was achieved through the Pdu pathway.

When glycerol induces the Pdu pathway, it is converted to 3-hydroxypropionaldehyde (3-HPA) by an AdoB_12_-dependent alcohol dehydratase comprised of PduCDE^12,18,19^. Under aerobic conditions, exogenous AdoB_12_ is required to initiate the conversion of glycerol to 3-HPA via the Pdu pathway^16^. 3-HPA is then converted by one of two aldehyde dehydrogenases, PduQ or PduP^12,19^. It is PduQ acting on 3-HPA in a NADH-dependent manner that produces 1,3-PDO; PduP produces 3-hydroxypropionyl-CoA in a NAD^+^/coenzyme A (CoA)-dependent manner, after which follows a further enzymatic cascade^12,31^. Key pathway enzymes and metabolites are illustrated in Figure 1A.

### S. enterica produces 1,3-PDO when pure or crude glycerol is the sole carbon and energy source

We sought to determine whether the addition of lab-grade glycerol and AdoB_12_ were sufficient to generate measurable quantities of 1,3-PDO in wild-type (WT) *S. enterica*. Over a 24-hour culture period, we detected both 3-HPA and 1,3-PDO in the WT sample starting around 12 h (Fig 1B). This point coincides with the complete consumption of glycerol, which starts to drop off after 8 h of incubation. At 12 h, 3-HPA peaks around 0.6 g/L. The highest concentrations of 1,3-PDO were not achieved until 12 h later, reaching approximately 0.8 g/L at 24 h. This titer corresponds to a yield of 0.16 g/g glycerol. Final biomass was approximately 1.4 g/L.

To establish whether the Pdu pathway is necessary for turning over glycerol to 1,3-PDO (via 3-HPA), we cultured a PocR-deficient strain (Δ*pocR)* so no pathway activation would occur, given the absence of the operon’s positive transcriptional regulator. Indeed, we observed no detectable levels of 3-HPA or 1,3-PDO over 24 h, even under the same media and culture conditions as the WT (Fig 1C). Glycerol alone is suitable for supporting growth in *S. enterica*, with Δ*pocR* biomass reaching approximately 1.8 g/L by 24 h. However, glycerol was not converted to 1,3-PDO without Pdu pathway activation.

As the enzymatic pathway branches after 3-HPA production, we hypothesized that 1,3-PDO production could be augmented by disrupting the competing pathway, catalyzed by PduP, and directing flux exclusively down the target branch, catalyzed by PduQ. Anticipating that we would achieve a higher conversion of glycerol to 1,3-PDO, we knocked out *pduP* and cultured the knockout strain (Δ*pduP*) under the same conditions as previously described (Fig 1D). However, by 24 h, 1,3-PDO ultimately reached comparable titers (0.7 vs. 0.8 g/L, *p* > 0.05) and yields (0.2 vs. 0.2 g 1,3-PDO/g glycerol, *p* > 0.05), as well as cell growth (1.3 vs. 1.4 g/L, *p* > 0.05), in the Δ*pduP* sample compared to WT.

Not only was there no increase in 1,3-PDO production in this strain, but it also showed a substantial accumulation of 3-HPA compared to WT. In the unmodified organism, 3-HPA peaks at 0.5 g/L at 12 h and shows a marked decrease to 0.2 g/L by 24 h. Without PduP, 3-HPA levels also reached a maximum at intermediate timepoints (0.8 g/L at 14 h) but at a 51% higher concentration than the WT sample (*p* < 0.05). Thereafter, 3-HPA remained elevated across the duration of the culture, with the final titer, 0.6 g/L, being 235% higher than in the WT culture (*p* < 0.005). Based on these results, we conclude that simply deleting *pduP* to eliminate the competing branch of the Pdu pathway is insufficient to drive flux toward 1,3-PDO and, in fact, causes an undesirable buildup of 3-HPA.

The industrial relevance of this pathway depends on whether crude glycerol is a viable substrate for growth and 1,3-PDO production. Thus, we set out to establish whether *S. enterica* could grow on crude glycerol and whether it would be converted to 1,3-PDO via the Pdu pathway (Fig 2A).

In fact, crude glycerol seems to be a slightly better carbon and energy source than pure glycerol, with WT cell growth exceeding 1.8 g/L after 24 h (35% increase, *p* < 0.01). Moreover, crude glycerol preserves 1,3-PDO production capabilities, with identical final titers and yields compared to cultures grown on pure glycerol (0.8 vs. 0.8 g/L and 0.16 vs. 0.16 g 1,3-PDO/g glycerol, respectively *p* > 0.5 in both instances). Interestingly, 3-HPA was only minimally detectable at one timepoint, reaching a maximum of 15 mg/L at 14 h.

Once again, in the *pocR* knockout sample, neither 1,3-PDO nor 3-HPA were detectable, while cell growth transpired and glycerol was completely consumed (Fig 2B). Since there was no 1,3-PDO in the absence of an active Pdu pathway, we conclude that Pdu enzymes facilitate crude glycerol conversion to 1,3-PDO.

### Final 1,3-PDO production can be improved with industrially relevant media composition changes

We also sought to identify how concentrations of our substrate and exogenous cofactor affected final 1,3-PDO production (Fig 3). From an industry perspective, high feedstock concentrations are necessary to maximize scalability^32^. Conversely, expensive cofactors should be minimized to reduce operating costs^33^.

We varied the starting concentration of glycerol to estimate the minimum feedstock dose and determine whether we could achieve continuous gains in 1,3-PDO by increasing feedstock dose alone (Fig 3A). When we used lab-grade glycerol as a substrate, some 1,3-PDO was detectable even at a dose of 1 g/L. A large increase in final 1,3-PDO titer was achieved when increasing starting dose from 1 to 2 g/L (209%, *p* < 0.005). However, the successive increase from 2 to 5 g/L led to a more modest increase in 1,3-PDO production (43%, *p* < 0.05). From 5 to 7 g/L, there was no statistical difference between final 1,3-PDO titers (*p* > 0.05); meanwhile, we observed a *decrease* in final 1,3-PDO titer when increasing the pure glycerol starting dose from 7 to 10 g/L (33%, *p* < 0.005).

Unlike pure glycerol, crude glycerol did not result in any detectable 1,3-PDO at 1 and 2 g/L doses. At 5 g/L, there was no difference in final 1,3-PDO production between pure and crude glycerol (*p* > 0.5). However, there was a sizeable increase in 1,3-PDO production when the starting dose of crude glycerol is raised from 5 to 7 g/L (94%, *p* << 0.005). Further increasing from 7 to 10 g/L crude glycerol does not result in a decrease in final 1,3-PDO, also unlike pure glycerol, although there was no significant difference between the final product titers (*p* > 0.1). Moreover, we measured significantly more 1,3-PDO when crude rather than pure glycerol was used at both the 7 and 10 g/L initial doses (increases of 72%, *p* < 0.005, and 180%, *p* < 0.01, respectively).

These results indicate that, while 1,3-PDO can be produced at lower starting doses of pure glycerol, higher titers are achievable under high feedstock doses. Additionally, using crude glycerol as a substrate delays the onset of diminishing marginal returns in 1,3-PDO as a function of starting dose. These findings have important scalability implications for using this pathway as a foundational bioprocess. Moreover, given the significant differences in performance across pure and crude glycerol, our results emphasize the need to test crude glycerol as early as possible in process development if a pathway or organism is proposed as a chassis for crude glycerol conversion. Relatively few studies have done this in the past^15,27,28,34^.

Understanding the impact of cofactors like AdoB_12_ on performance is also critical for industrial relevance. Not only is AdoB_12_ expensive, but using less than needed to achieve saturation would decrease the rate of turnover of glycerol to 3-HPA. This could further benefit the glycerol-to-1,3-PDO conversion process by reducing the effective concentration of 3-HPA within cells, protecting them from its toxic effects and preserving their fitness as biological chasses. Hypothesizing that AdoB_12_ concentrations below saturation would improve final 1,3-PDO titers, we tested the effect of saturating (150 nM) and limiting (20 nM) AdoB_12_, as well as one midpoint concentration (85 nM), on 24 h 1,3-PDO production (Fig 3B).

Under aerobic conditions, AdoB_12_ must be added. At 0 nM, no 1,3-PDO is produced from glycerol. Of the three non-zero AdoB_12_ conditions we tested, the limiting concentration proved to result in the highest final 1-3-PDO titer. We detected no significant difference between the saturating and midpoint concentrations (0.47 vs. 0.48 g/L, *p* > 0.5), but using a limiting quantity of AdoB_12_ yielded an increase in 1,3-PDO production of 73% compared to the saturating concentration (*p* < 0.05). Moreover, using the limiting concentration reduced peak 3-HPA concentration by 95% compared to both higher AdoB_12_ concentrations (p < 0.05) and resulted in no residual 3-HPA at 24 h. Limiting AdoB_12_ did not negatively impact cell growth or glycerol consumption (complete growth curves in Fig S2). It may thus be feasible to minimize AdoB_12_ addition during bioprocess scale-up for this pathway.

## Conclusion

*S. enterica* is fast-growing, genetically tractable, and well characterized and offers several additional advantageous characteristics that make it a strong potential platform for 1,3-PDO production from crude glycerol. It has a native glycerol assimilation mechanism, so it can utilize crude glycerol as a sole carbon and energy source while other proposed chasses cannot^13,35–37^. Additionally, we showed here that 1,3-PDO production can occur aerobically in *Salmonella*.

Thus far, much of the research around biological conversion of glycerol to 1,3-PDO centers around the pathway governed by the *dha* (dihydroxyacetone) regulon, whose enzymes are oxygen sensitive even in facultative organisms (many 1,3-PDO-producing organisms are obligate anaerobes)^38–40^. Furthermore, Dha enzymes are readily inactivated by glycerol and 3-HPA^40–43^. *S. enterica* offers its Pdu pathway as an alternative route for generating 1,3-PDO.

Engineering Pdu pathway enzymes holds great promise for improving on the performance reported in this work. Increasing enzymes’ affinity for glycerol and 3-HPA and their rate of 3-HPA-to-1,3-PDO turnover could both lead to substantial gains in 1,3-PDO yield. Moreover, the Pdu microcompartment (MCP) structure could be engineered in tandem to the pathway to the advantage of this conversion. Chimeric shell compositions and alterations to pore sites have been already utilized to enhance pathway performance in other MCP research^44,45^.

In this work, we have demonstrated that *S. enterica* can convert glycerol, including crude glycerol, to 1,3-PDO given a minimal media formulation. Compared to other studies in which greater levels of iteration were conducted,^15,27,28,34^ our titers and yields may appear low. However, our study is a foundational first step toward continuous development of the *S. enterica* and the Pdu pathway as tools for bioconversion.

## Acknowledgements

Edward Y. Kim and Christopher M. Jakobson generated our knockout strains. We greatly appreciate the generous support of Zach Waickman and the Loyola Biodiesel Lab, who donated all crude glycerol used in this project.

## Author contributions

**Madeline Joseph**: conceptualization, formal analysis, investigation, visualization, writing (original draft). **Brett Palmero**: conceptualization, methodology, writing (reviewing and editing). **Nolan Kennedy**: conceptualization, project management, supervision, writing (reviewing and editing). **Danielle Tullman-Ercek**: conceptualization, funding acquisition, project management, resources, supervision, writing (reviewing and editing).

## Declarations

The authors declare no conflicts of interest.

