## Supplemental Text for "Glycerol alone effects 1,3-propanediol production via the aerobic propanediol utilization pathway in *Salmonella enterica*"

### Materials and Methods

*Determination of crude glycerol purity*

The exact composition of our crude glycerol sample was unknown. We used high performance liquid chromatography (HPLC) to determine its glycerol content. First, we diluted a stock solution comprised of 50 g of crude glycerol to 1 L of water (5% w/v) by a factor of 10 to achieve a 0.5% w/v composition of *crude* glycerol (our standard dose for pure glycerol-fed cultures). Then, we ran this sample on HPLC against a pure glycerol standard. The measured glycerol concentration was approximately 0.25% w/v, so the 1 L stock solution, which contained 50 g of *crude* glycerol, was 2.5% w/v glycerol. This corresponds to a 50% glycerol purity by weight for our crude sample. We used this value to adjust the feedstock volume we added when growing cultures on crude glycerol.

*Biomass measurement conversions*

When we determined cell density using $OD_{600}$, we converted these measurements to dry cell mass (DCM), in g/L, using linear standard curves (Fig S1)[1]. We calculated the conversion factor, in g/L per unit of $OD_{600}$, from a linear correlation curve composed of 5 pairs of DCM-$OD_{600}$ data. All media and culture conditions were the same as described in the main article, except these cultures grew in 3 L glass Erlenmeyer flasks with a working volume of 500 mL. We obtained cell mass measurement correlations for all strains of *S. enterica* tested in this work: wild-type (WT), a *pocR* knockout (Δ*pocR*), and a *pduP* knockout (Δ*pduP*). For WT, Δ*pocR*, and Δ*pduP*, the linear conversion factors were 0.4201 ($R^2$ = 0.993), 0.4415 ($R^2$ = 0.981), and 0.4148 ($R^2$ = 0.995) g/L per unit of $OD_{600}$, respectively.

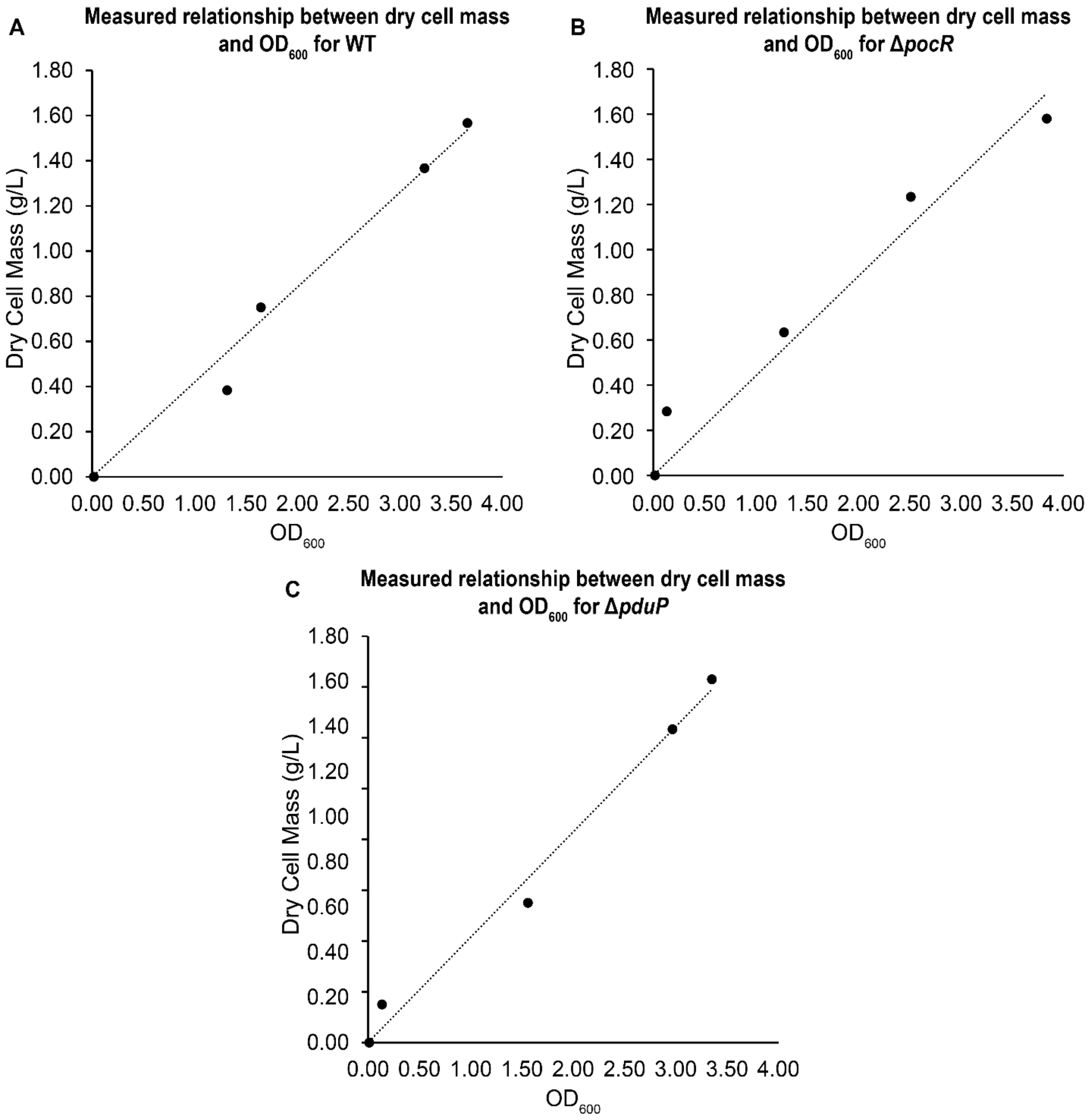


**Figure S1.** Linear relationship between experimentally measured $OD_{600}$ and dry cell mass (g/L) for **A)** wild-type (WT), **B)** PocR-deficient (Δ*pocR*), and **C)** PduP-deficient (Δ*pduP*) strains of *Salmonella enterica.*

*Statistics*

As all experiments were performed in biological triplicate, sample size (n) is 3 in all cases. Standard deviation is used to denote variation within samples. When comparing values, means and standard deviations for these triplicate data were used. We calculated the significance of these comparisons as *p*-values using Welch's Two-Sample *t*-test for test statistics and the Welch-Satterthwaite formula for degrees of freedom. We report values, standard deviations, *t*-statistics, degrees of freedom, and *p*-values for data referenced in the article and supplement in Supplementary Table 1.

**Table S1.** Calculated averages, standard deviations, percent-changes (compared to the baseline value), *t*-statistics, degrees of freedom (df), and *p*-values for select data comparisons. Sample size is 3 in all cases.

### Results

*Limiting $AdoB_{12}$ does not negatively affect cell growth or glycerol consumption*

We collected time-resolved cell growth and metabolite data in WT for different doses of exogenous $AdoB_{12}$ (Fig S2). By 24 h, there was no significant difference in the biomass accumulation between saturating or midpoint $AdoB_{12}$ and limiting $AdoB_{12}$ (0.9 g/L vs. 1.2 g/L, $p >$ 0.05 in both cases). Moreover, all cultures had the same characteristic glycerol consumption pattern: glycerol falls off rapidly at 12 h and reaches >80% consumption by 16 h. Thus, minimizing $AdoB_{12}$ does not seem to detrimentally alter the key industrially relevant behaviors of cells in the context of glycerol-to-1,3-PDO production.

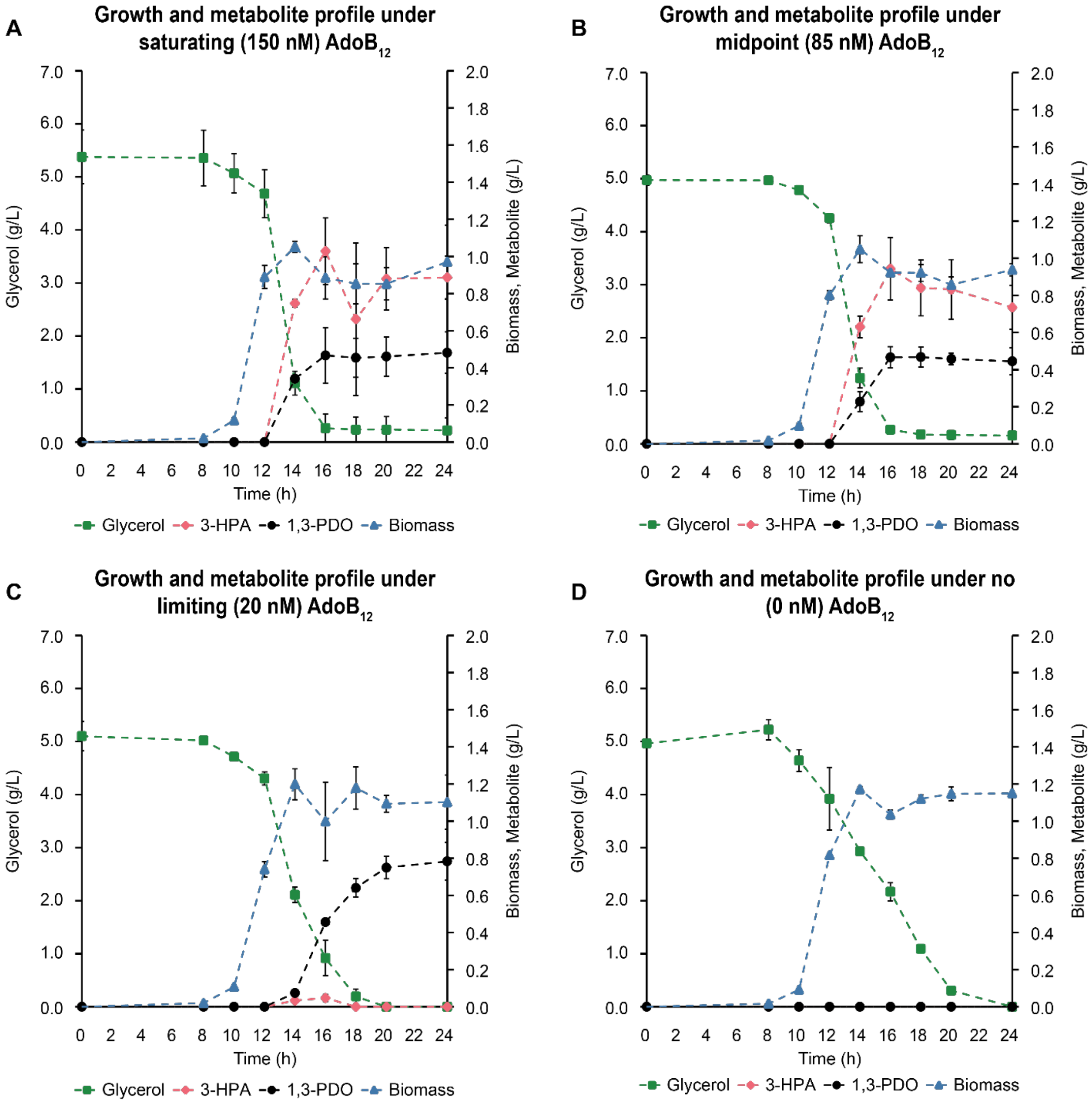


**Figure S2.** In wild-type *S. enterica*, time courses for varying doses of $AdoB_{12}$: **A)** saturating (150 nM), **B)** midpoint (85 nM), **C)** limiting (20 nM), and **D)** none (0 nM).
